## Supporting Information for "Nanomechanics and co-transcriptional folding of Spinach and Mango"

[1]*Department of Materials Science and Engineering, University of Illinois at Urbana-Champaign, Urbana IL, USA;* [2]*Department of Biophysics and Biophysical Chemistry, Johns Hopkins University, Baltimore, MD, USA;* [3]*Department of Biophysics, Johns Hopkins University, Baltimore, MD, USA;* [4]*Department of Biomedical Engineering, Johns Hopkins University, Baltimore, MD, USA;* [5]*Howard Hughes Medical Institute, USA*

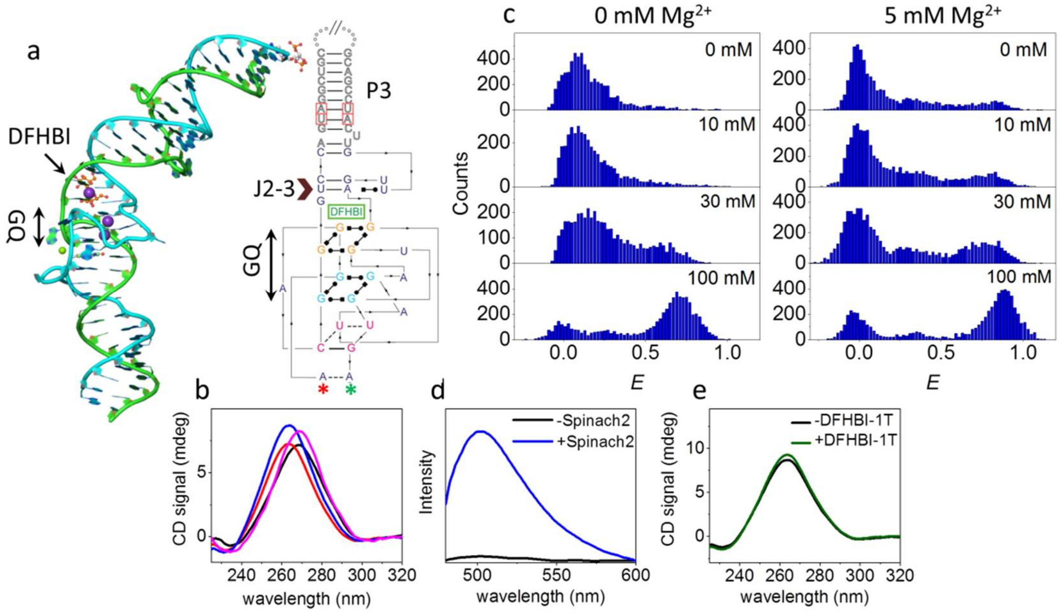


**Figure S1.** (a) Co-crystal structure of Spinach2 and DFHBI.[1] DFHBI is shown in orange and the purple balls represent intercalating ions (left). The $U_{32}$-$A_{64}$-$U_{61}$ base triple, forming the J2-3 junction between GQ and the flanking duplex stem (P3) is shown by a brown arrowhead. The relative positions of Cy5 and Cy3 in the Spinach2 construct are indicated by red and green asterisks respectively. (b) CD spectra of Spinach2 in 0 mM $K^+$ (black), 100 mM $K^+$ (red), 5 mM $Mg^{2+}$ (magenta) and 100 mM $K^+$ and 5 mM $Mg^{2+}$ (blue) containing buffers. (c) *E* histogram of Spinach2 with varying concentrations of $K^+$, with (right) and without (left) $Mg^{2+}$, in the absence of force. (d) Emission spectra of DFHBI-1T in the Spinach bound and unbound states ($\lambda_{exc}$ = 460 nm). (e) CD spectra of Spinach2 with and without DFHBI-1T.

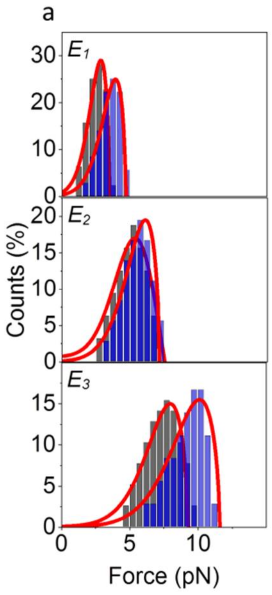


b

| | | $\Delta x_u^‡$ (nm) | $\Delta G^‡$ ($k_BT$) | $\tau_u(0)$ (s) |
|---|---|---|---|---|
| -DFHBI-1T | $E_1$ | 10.1±1.2 | 5.8±0.7 | 30.2±6.1 |
| | $E_2$ | 6.8±0.8 | 7.3±1.3 | 202.5±29.2 |
| | $E_3$ | 5.8±0.8 | 8.6±1.0 | 246.5±41.7 |
| +DFHBI-1T | $E_1$ | 8.0±0.9 | 6.6±1.6 | 103.7±23.4 |
| | $E_2$ | 6.7±0.8 | 7.7±1.4 | 270.6±38.3 |
| | $E_3$ | 4.2±0.7 | 9.2±2.2 | 890.7±230.8 |

**Figure S2.** (a) Distributions of unfolding forces corresponding to the $E_1$, $E_2$ and $E_3$ states of Spinach2 with (blue) and without (black) DFHBI-1T. The red curves represent unfolding force distributions predicted from the Dudko-Szabo model.[2,3] The free energy parameters are tabulated in (b). The errors are calculated from 95 % confidence intervals.

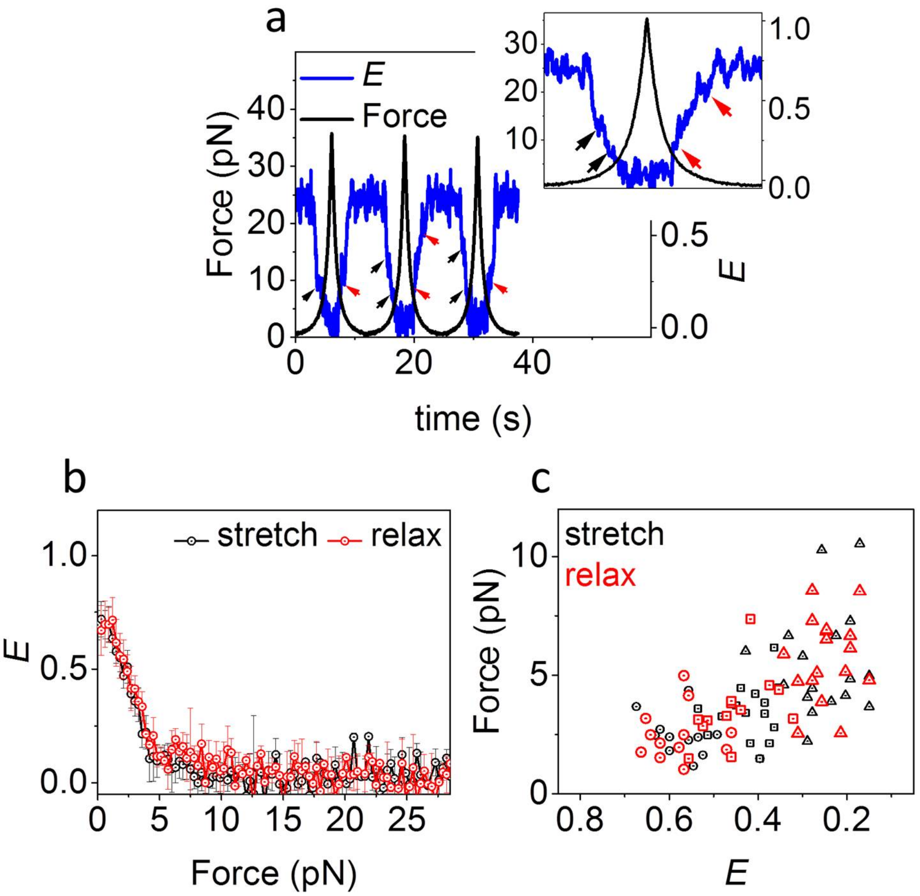


**Figure S3.** (a) smFRET time trajectory (20 ms integration time) of Spinach2 in 100 mM $K^+$ over three pulling cycles. A blown-up image of cycle 2 is shown in the inset. The black and red arrows indicate unfolding and refolding steps respectively. (b) Average *E* vs force response of Spinach2 (*N*=36). (c) Force vs *E* corresponding to each unfolding (stretch) and refolding (relax) step.

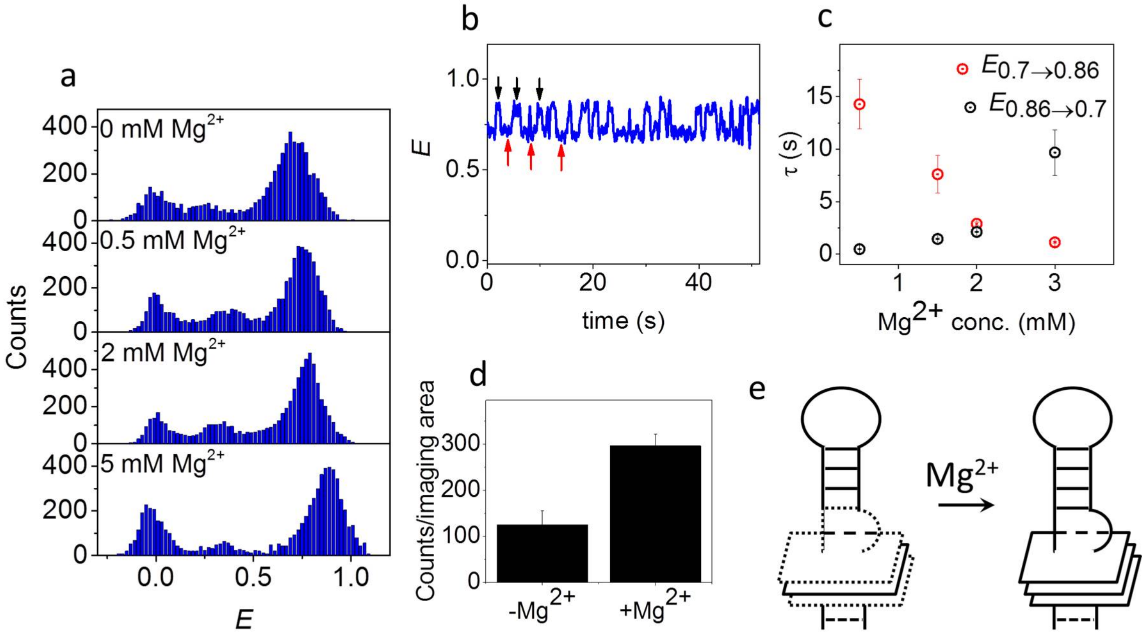


**Figure S4.** (a) $E$ histograms of Spinach2 in buffer containing 100 mM $K^+$ and varying concentrations of $Mg^{2+}$, in the absence of force. (b) smFRET time trajectory of Spinach2 (30 ms integration time) in 100 mM $K^+$+2 mM $Mg^{2+}$ buffer, in the absence of force. The red and black arrows indicate transitions between the $E_{0.7}$ and $E_{0.86}$ states. (c) Dwell times of $E_{0.7}$ and $E_{0.86}$ states in 100 mM $K^+$, with varying $Mg^{2+}$ concentrations. (d) DFHBI-1T binding to Spinach2 in 100 mM $K^+$, with and without $Mg^{2+}$. (e) A schematic showing probable stabilization of the stem loop-junction of Spinach2 GQ with $Mg^{2+}$.

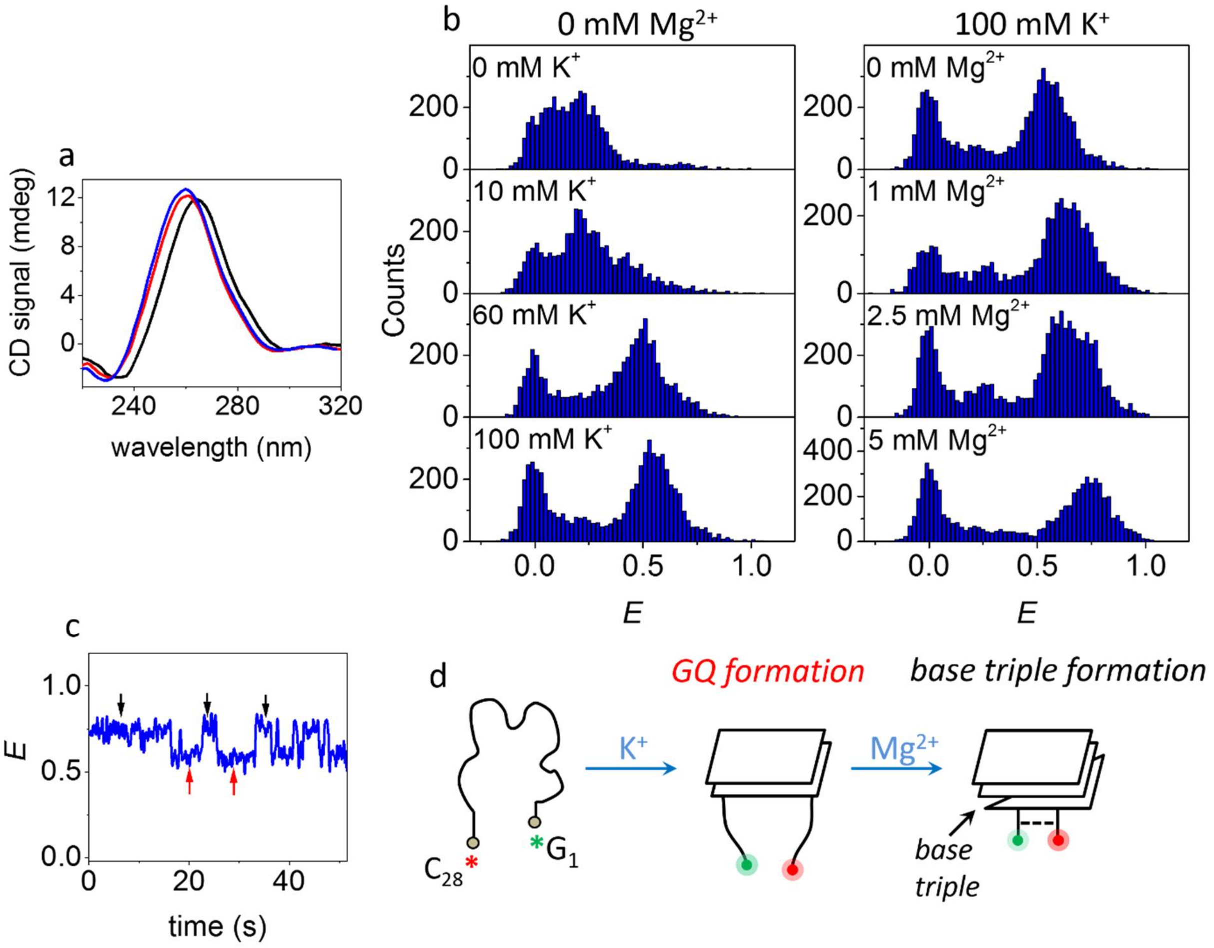


**Figure S5.** (a) CD spectra of *i*MangoIII in 0 mM $K^+$ (black), 100 mM $K^+$ (red) and 100 mM $K^+$ and 5 mM $Mg^{2+}$ (blue) containing buffers. (b) *E* histogram of *i*MangoIII with varying concentrations of $K^+$ (left) and $Mg^{2+}$ (right) in the absence of force. (c) smFRET time trajectory of *i*MangoIII (30 ms integration time) in 100 mM $K^+$+2.5 mM $Mg^{2+}$ buffer, in the absence of force. The red and black arrows indicate transitions between two *E* states. (d) Mechanism of ion-induced *i*MangoIII folding suggested with reference to (c). The green and red asterisks represent the Cy3 and Cy5 dyes, adjacent to $G_1$ and $C_{28}$ respectively.

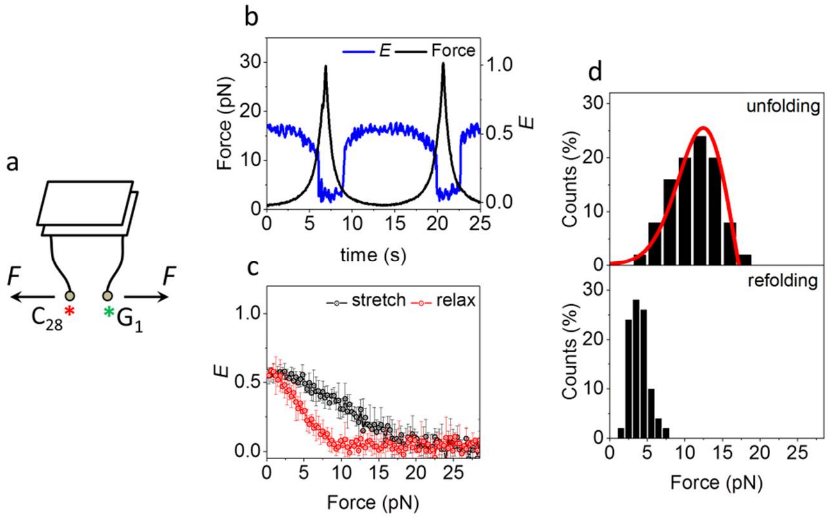


**Figure S6.** (a) A schematic of *i*MangoIII in 100 mM $K^+$, under tension. The green and red asterisks represent the Cy3 and Cy5 dyes, adjacent to $G_1$ and $C_{28}$ respectively. (b) A representative smFRET trajectory of *i*MangoIII in 100 mM $K^+$, over two pulling cycles. (c) Average $E$ vs force response. (d) Distributions of unfolding and refolding forces ($N$=50 for both). The red curves represent force distributions estimated from the Dudko-Szabo model[2,3].

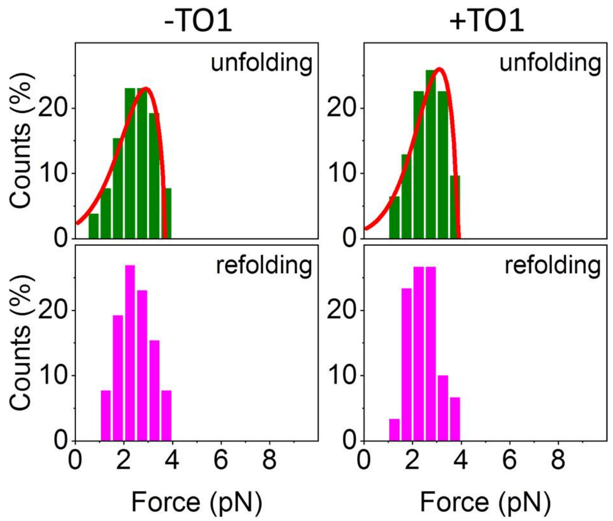


**Figure S7.** Distributions of unfolding and refolding forces corresponding to transitions between $E_{0.74}$ and $E_{0.6}$ in *i*MangoIII. Data were acquired in buffer containing 100 mM $K^+$ and 5 mM $Mg^{2+}$, with (right, $N$=50) and without (left, $N$=62) TO1. The red curves represent force distributions estimated from the Dudko-Szabo model[2,3]. The free energy parameters used are: $\Delta x_u^{\ddagger}$ 7.2±1.8 nm (left) and 6.8±1.8 nm (right), $\Delta G^{\ddagger}$ 4.4±1.0 $k_BT$ (left) and 4.2±1.0 $k_BT$ (right) and $\tau_u(0)$ 50.4±11.3 s (left) and 87.1±22.1 s (right). The errors are calculated from 95 % confidence intervals.

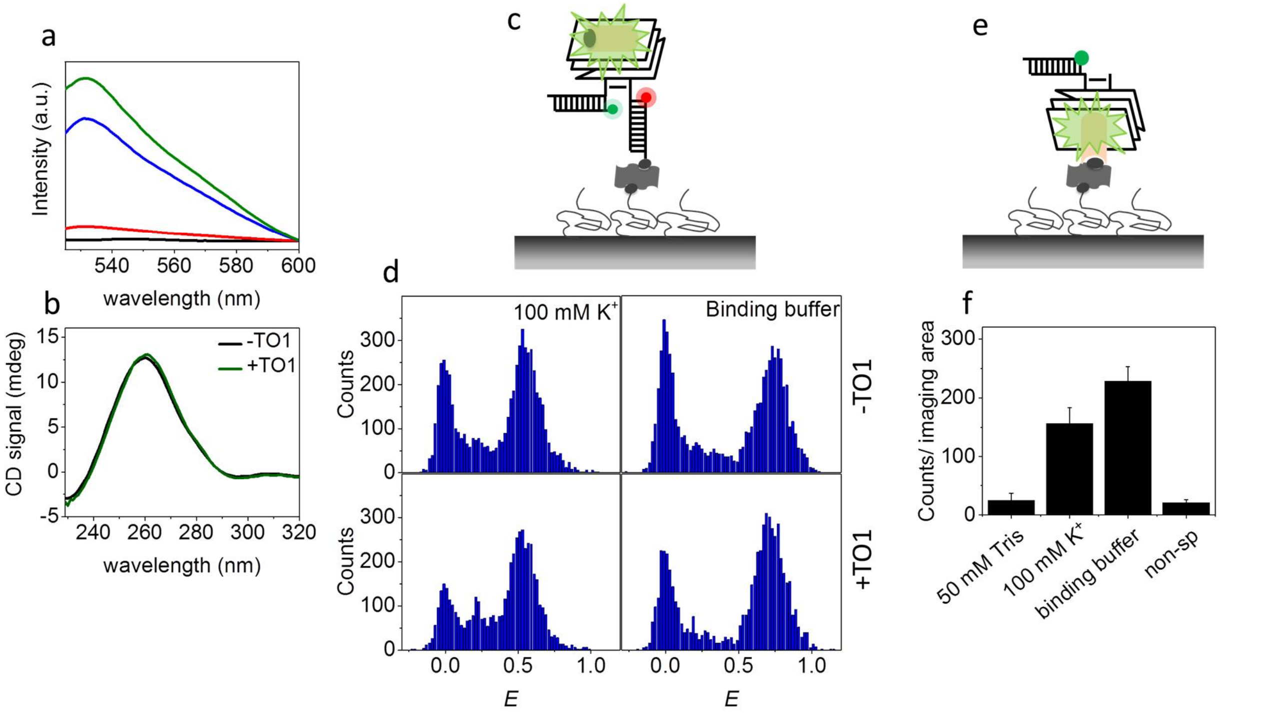


**Figure S8.** (a) Emission spectra ($\lambda_{exc}$ = 485 nm) of TO1-*i*MangoIII complex in 50 mM Tris pH 7.5 only (red), 100 mM $K^+$ (blue) and 100 mM $K^+$ and 5 mM $Mg^{2+}$ (green). Black curve represents emission spectrum of TO1 only. (b) CD spectra of *i*MangoIII with and without TO1. (c) Schematic representation of TO1 (in black and orange) bound to *i*MangoIII on a single molecule platform. (d) *E* histograms of *i*MangoIII in the presence and absence of TO1. (e) TO1 was pulled down on the single molecule surface via biotin-neutravidin linkage. Aptamer binding was visualized with Cy3-labelled *i*MangoIII. (f) Quantification of *i*MangoIII bound TO1, immobilized on the PEG-passivated surface. All measurements were done in binding buffer containing 100 mM $K^+$ and 5 mM $Mg^{2+}$, unless otherwise specified.

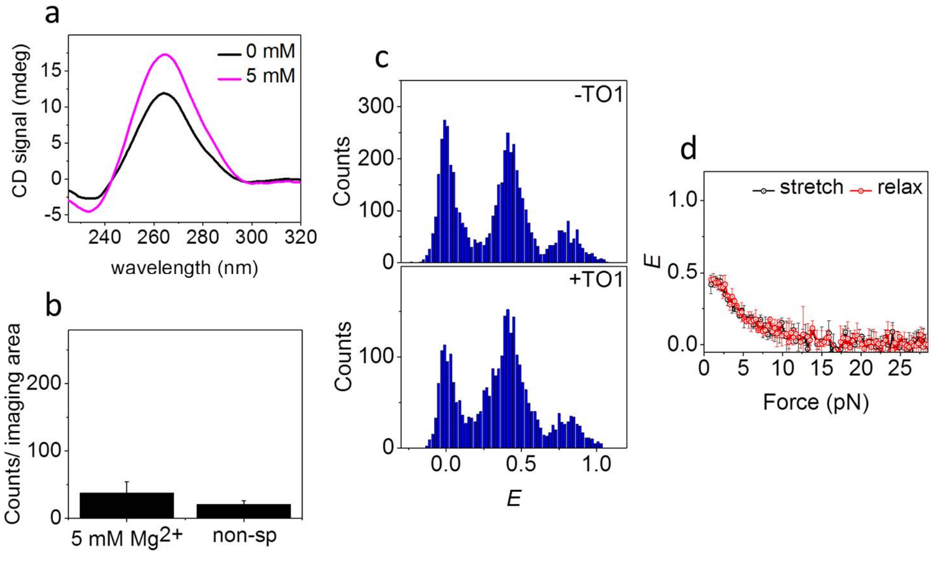


**Figure S9.** (a) CD spectra of *i*MangoIII in 0 mM (black) and 5 mM $Mg^+$ (magenta). (b) Quantification of *i*MangoIII binding to TO1, immobilized on the single molecule surface, in 5 mM $Mg^{2+}$. (d) *E*

histogram of *i*MangoIII with (bottom) and without (top) TO1 in 5 mM $Mg^{2+}$ in the absence of force. (e) *E* vs force response of the dominant mid-*E* population of *i*MangoIII in 5 mM $Mg^{2+}$.

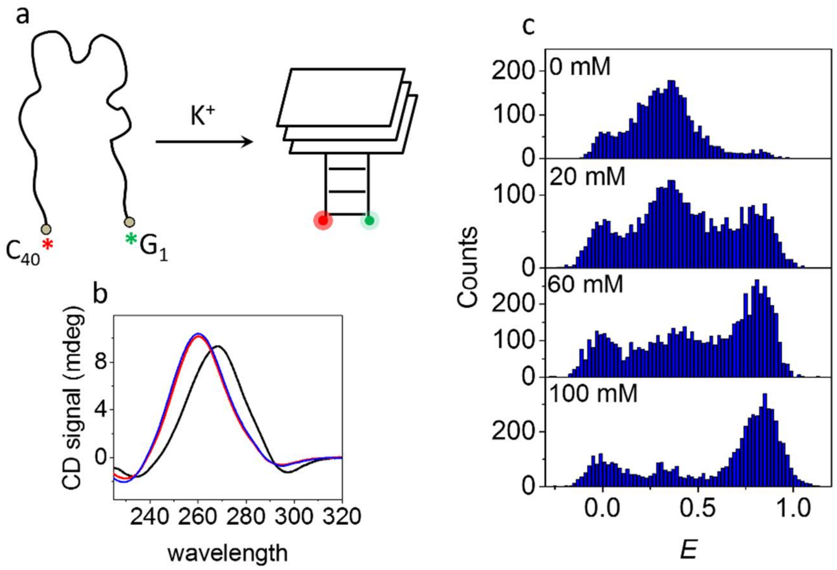


**Figure S10.** (a) $K^+$ induced folding of MangoIV. The green and red asterisks represent the Cy3 and Cy5 dyes, adjacent to $G_1$ and $C_{40}$ respectively. (b) CD spectra of MangoIV in 0 mM $K^+$ (black), 100 mM $K^+$ (red) and 100 mM $K^+$+5 mM $Mg^{2+}$ (blue). (c) *E* histograms of MangoIV under varying concentrations of $K^+$, in the absence of force.

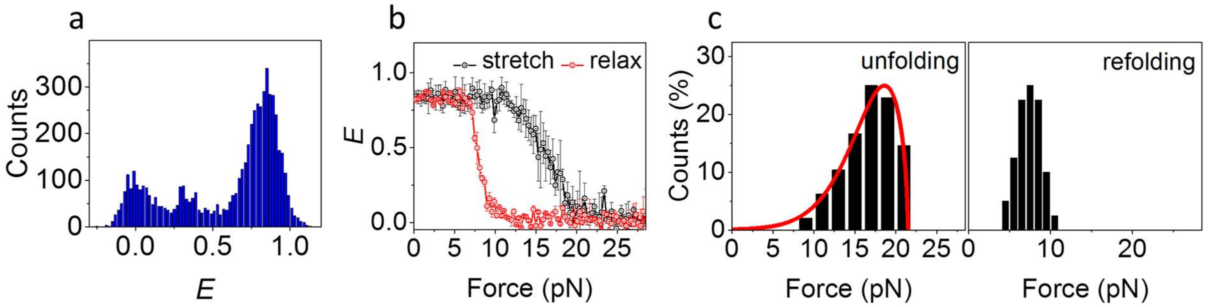


**Figure S11.** (a) *E* histogram of MangoIV, in the absence of force. (b) Average *E* vs force response. (c) Distributions of unfolding (left) and refolding forces (right). (*N*=48 for both). The red curves represent force distributions estimated from the Dudko-Szabo model (left)[2,3]. All measurements were performed in a buffer containing 100 mM $K^+$.

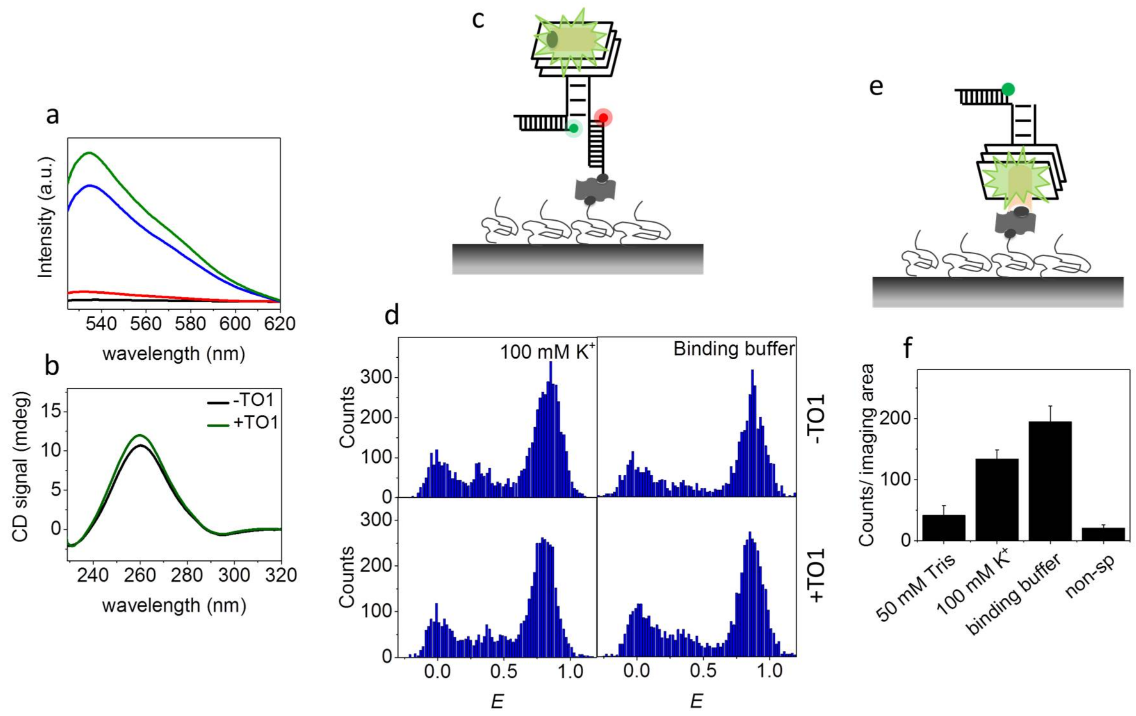


**Figure S12.** (a) Emission spectra ($\lambda_{exc}$ = 485 nm) of TO1-MangoIV complex in 50 mM Tris pH 7.5 only (red), 100 mM $K^+$ (blue) and 100 mM $K^+$ and 1 mM $Mg^{2+}$ (green). Black curve indicates emission spectrum of TO1 only. (e) CD spectra of MangoIV with and without TO1. (c) Schematic representation of TO1 (in black and orange) bound to MangoIV on the single molecule platform. (d) *E* histograms of MangoIV in the presence and absence of TO1. (e) Binding of Cy3-labelled MangoIV to TO1, immobilized on the single molecule surface via biotin-neutravidin linkage. (f) Quantification of MangoIV bound TO1 in (e).

| | | $\Delta x_u^{\ddagger}$ (nm) | $\Delta G^{\ddagger}$ ($k_BT$) | $\tau_u(0)$ (s) |
|---|---|---|---|---|
| *i*MangoIII | Buffer 1 | 4.2±0.5 | 8.8±1.4 | 798±190 |
| | Buffer 2 | 3.8±0.7 | 8.7±1.4 | 1412±289 |
| | Buffer 3 | 2.8±0.5 | 8.2±1.2 | 566±134 |
| MangoIV | Buffer 1 | 2.8±0.6 | 9.7±1.8 | 1726±364 |
| | Buffer 2 | 3.0±0.3 | 9.9±0.9 | 1191±101 |
| | Buffer 3 | 2.2±0.5 | 8.5±1.9 | 389±87 |

**Figure S13.** The free energy parameters of *i*MangoIII and MangoIV. The measurements were done in buffer containing 100 mM $K^+$ and 5 mM $Mg^{2+}$. (c) Comparison of the unfolding free energy parameters of the Mango aptamers.[2,3] Buffer 1: 100 mM $K^+$, Buffer 2: fluorogen binding buffer, 100 mM $K^+$ and 5 mM $Mg^{2+}$ (*i*MangoIII) or 1 mM $Mg^{2+}$ (MangoIV) and Buffer 3: TO1 in fluorogen-binding buffer. The errors are calculated from 95 % confidence intervals.

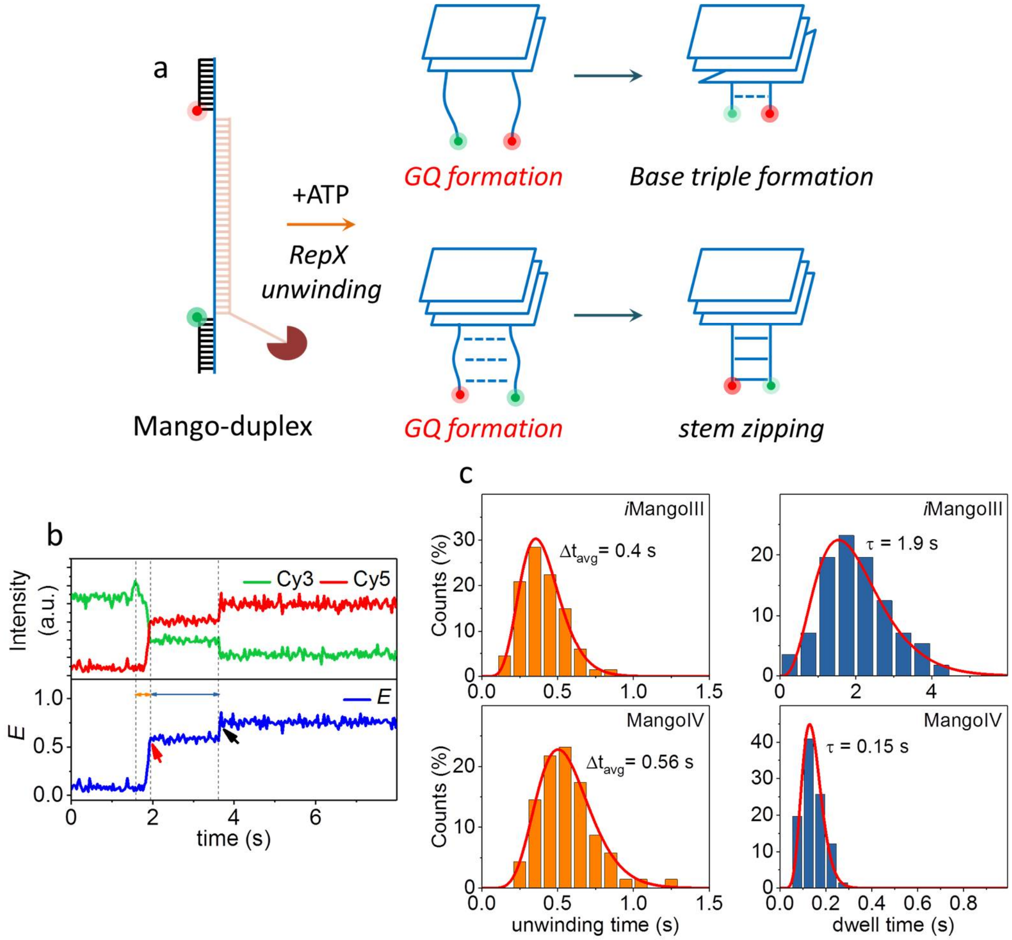


**Figure S14.** (a) Schematic showing possible mechanisms of vectorial folding of *i*MangoIII (top) and MangoIV (bottom). (b) Representative smFRET time trajectory of *i*MangoIII. Rep-X translocation towards the *i*MangoIII duplex is reflected by protein induced fluorescence enhancement (PIFE, black arrow head) and unwinding time of duplex was calculated using the PIFE peak as the starting point. The time taken for unwinding and GQ formation is shown by the orange arrow. Time lag between GQ and base triple formation is shown by the blue arrow. (c) Histograms of time taken for duplex-unwinding/GQ formation (left) and dwell time of GQ-only intermediate prior to base triple formation (*i*MangoIII) or stem zipping (MangoIV) (right). The average unwinding times and dwell times were calculated from fitting to gamma distributions (red curves).

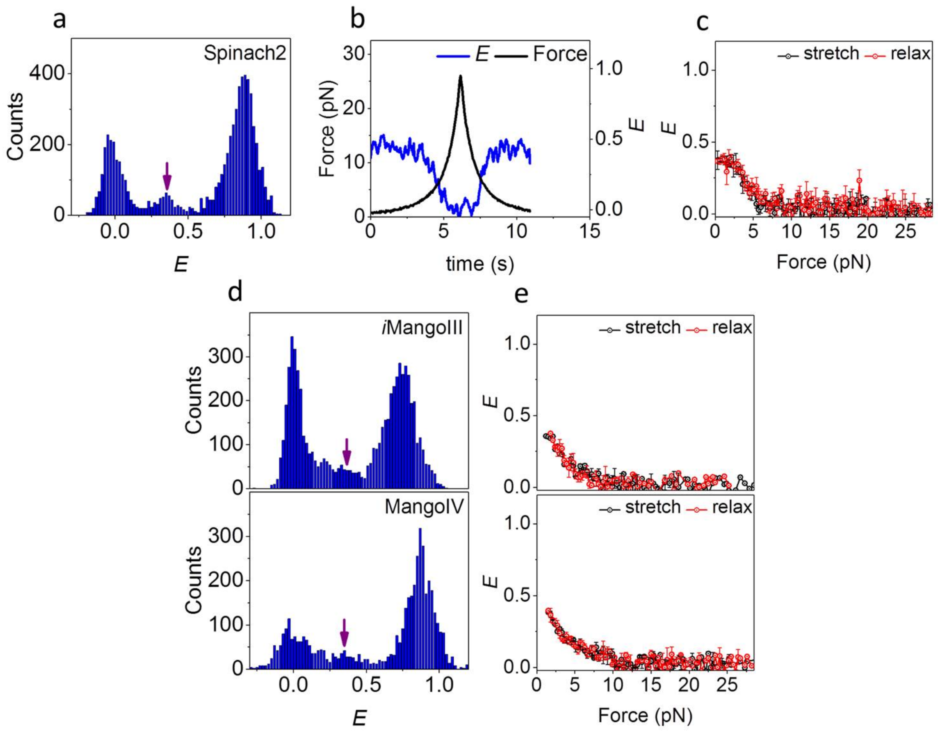


**Figure S15.** (a) *E* histogram of Spinach2 in the absence of force. The purple arrow indicates a mid-*E* population. (b) A representative single molecule time trajectory in the mid-*E* state (20 ms integration time). (c) Average *E* vs force response of the population indicated in (a). (d) *E* histograms of *i*MangoIII (top) and MangoIV (bottom) in the absence of force. (e) Average *E* vs force responses of mid-*E* *i*MangoIII (top) and MangoIV (bottom). All measurements were performed in Rep-X unwinding buffer conditions.

All error bars represent standard errors.

### References


1 Warner, K. D. *et al.* Structural basis for activity of highly efficient RNA mimics of green fluorescent protein. *Nat. Struct. Mol. Biol.* **21**, 658-663 (2014).

2 Dudko, O. K., Hummer, G. & Szabo, A. Intrinsic rates and activation free energies from single-molecule pulling experiments. *Phys. Rev. Lett.* **96**, 108101 (2006).

3 Dudko, O. K., Hummer, G. & Szabo, A. Theory, analysis, and interpretation of single-molecule force spectroscopy experiments. *Proc. Natl. Acad. Sci. U S A.* **105**, 15755-15760 (2008).
